## Supplemental Information for "Epistatic models predict mutable sites in SARS-CoV-2 proteins and epitopes"


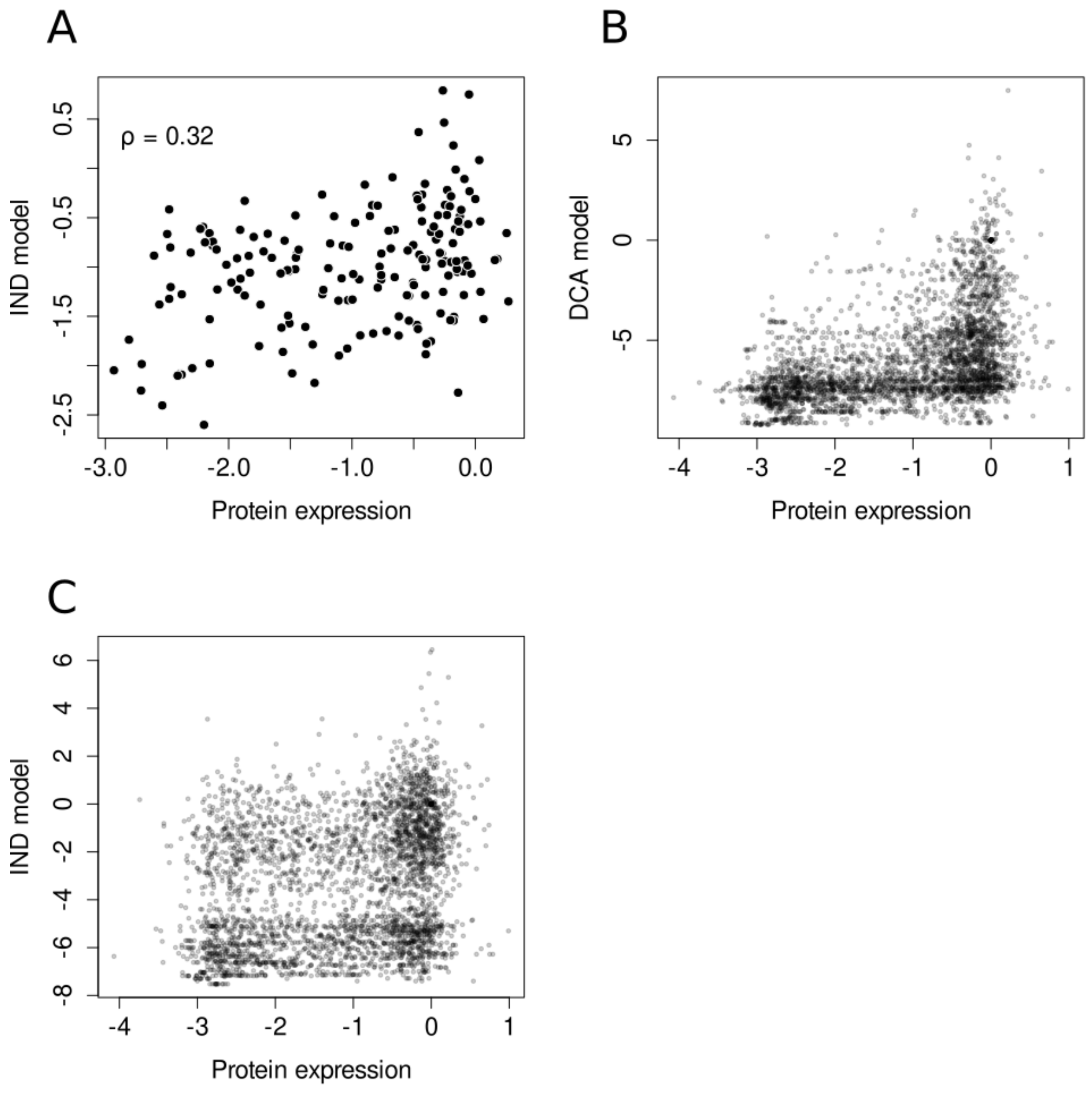


*Fig. S1 A) Experimental protein expression for the 178 positions of the RBD as a function of the predicted effect by the IND model. The effects of 3355 single mutations in the 178 positions of the RBD measured by the experimental protein expression in the x-axis and predicted by the DCA model (B) and the IND model (C) in the y-axis.*

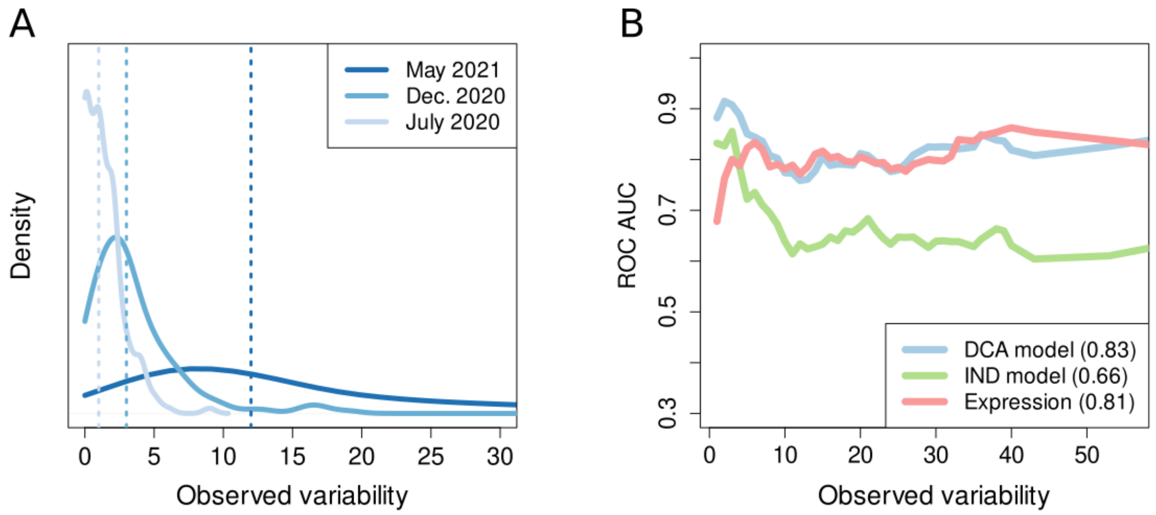


*Fig. S2. A) Distributions of observed variability using the genomes available at July 2020, December 2020 and May 2021. The vertical dashed lines represent the median of each distribution and is used as a cutoff to distinguish between low- and high-variability positions. B) AUC for ROC curves using cutoffs of variability in the interval [1,56]. The mean AUC is shown in the legend.*

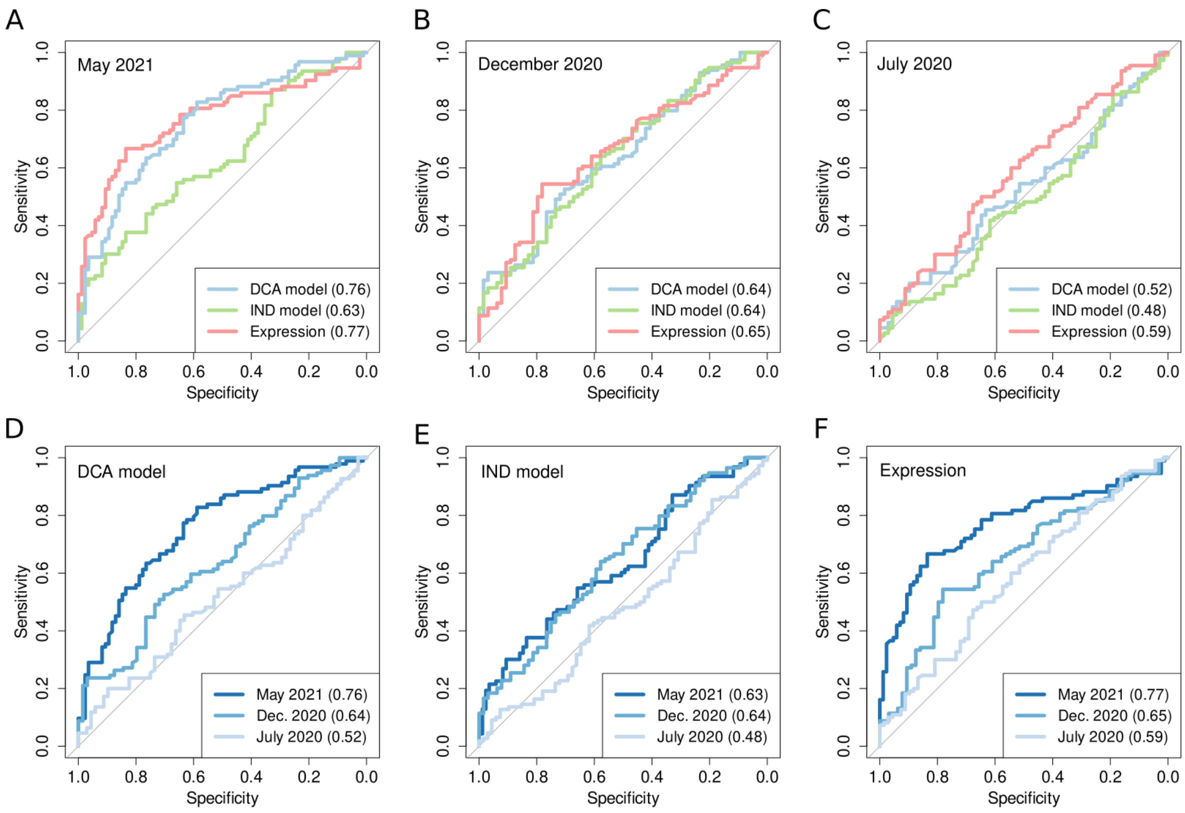


*Fig S3. ROC curves for positions with low versus high observed variability for the 3 predictors with observed variability derived from all the SARS-CoV-2 genomes available at May 2021 (A), December 2020 (B), and July 2020 (C). ROC curves for positions with low versus high observed variability, where the observed variability is quantified with the SARS-CoV-2 genomes available at July 2020, December 2020, and May 2021 for the prediction coming from the DCA model (D), IND model (E), and protein expression (F).*

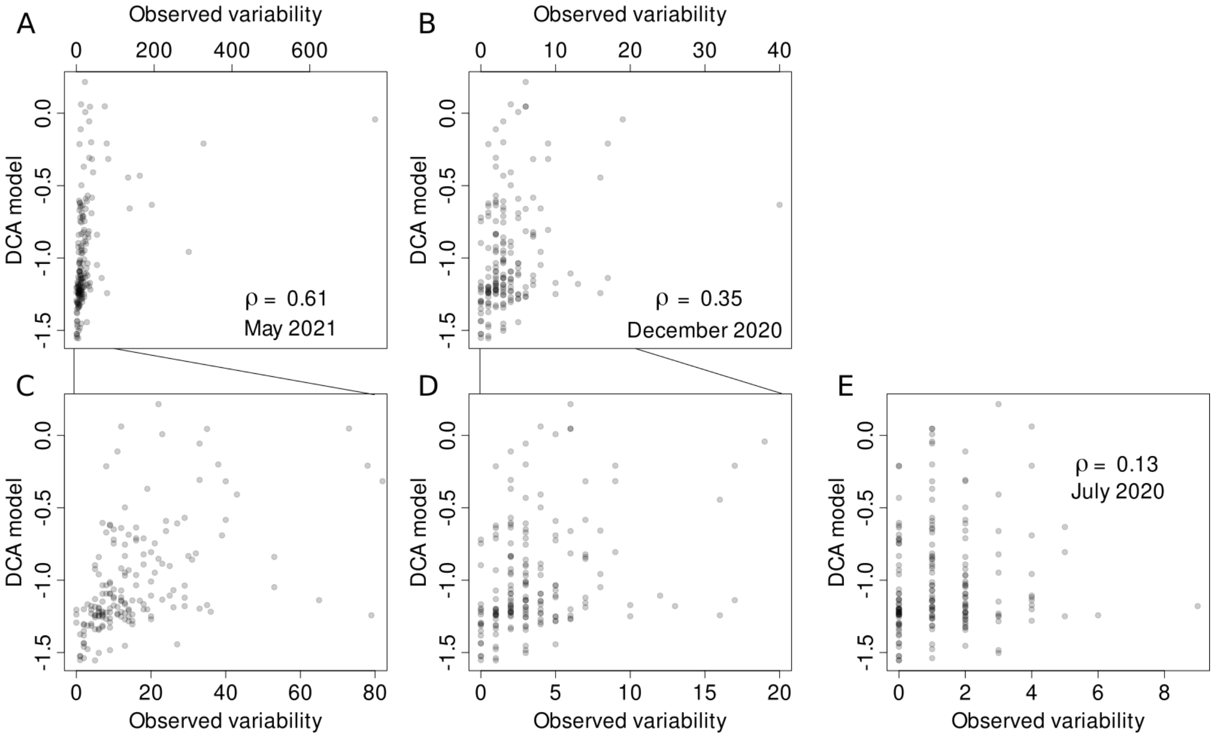


*Fig. S4. Observed variability (estimated with GISAID data till May 2021, panels A and C; December 2020, B and D; July 2020, E) compared to the DCA predictions. The whole range of observed variabilities is shown in panels A, B, and E. In panels C and D, a more restricted interval of observed variabilities is shown to improve the visibility.*

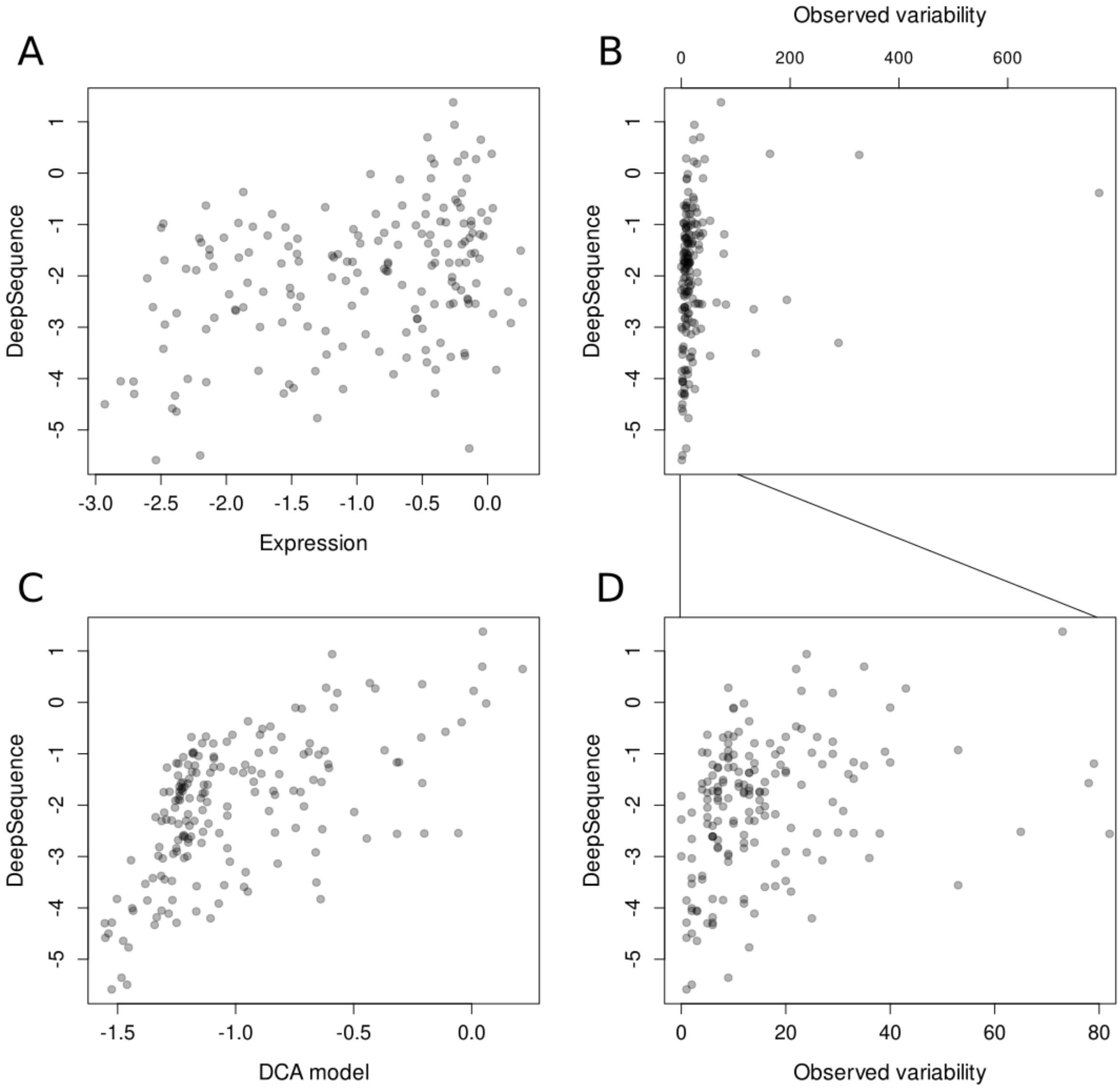


*Fig. S5. A) Scatter plot between protein expression and DeepSequence scores. Comparison between observed variability and DeepSequence scores for the whole range of observed variability in May 2021 (B) and in the interval of [0,80] (D). C) DeepSequence scores as a function of the DCA model scores.*

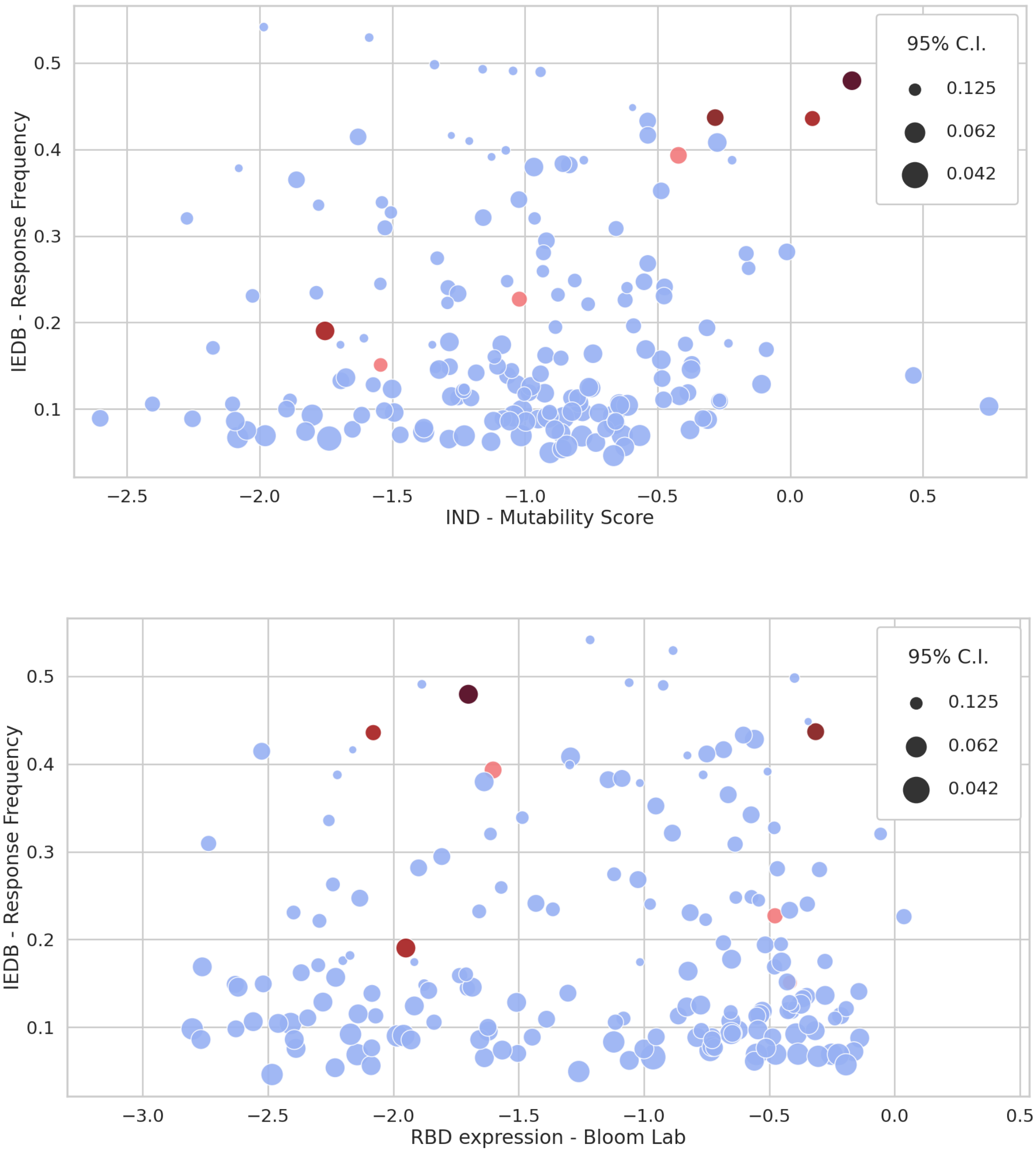


*Fig. S6. The IEDB-Response Frequency as a function of the IND mutability score (upper panel) or protein expression (lower panel) for each position of the RBD domain. The enrichment of VOC/VOI mutations becomes less pronounced as compared to the DCA score.*

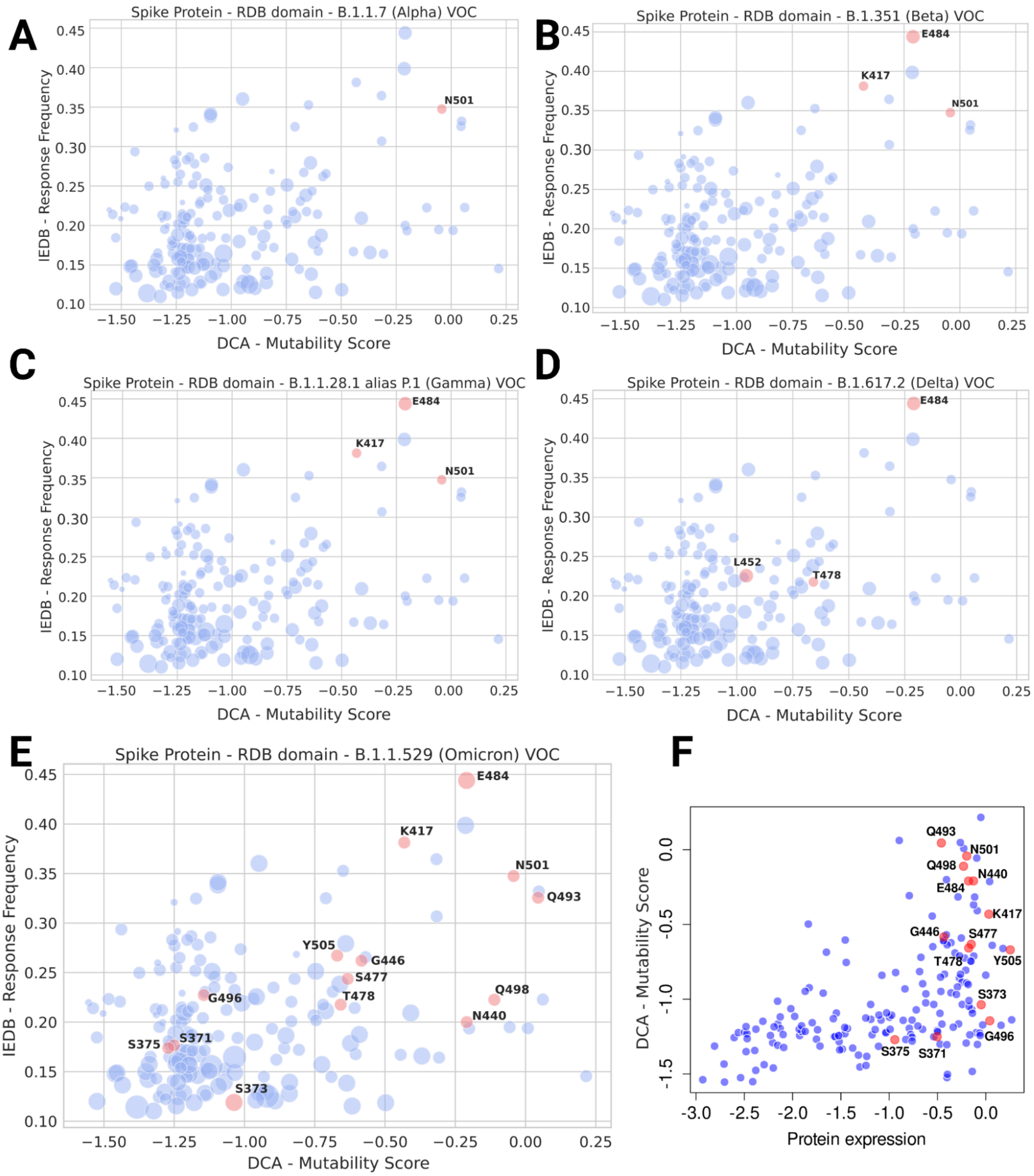


*Fig. S7 - The IEDB-Response Frequency versus the DCA mutability score with updated IEDB response frequencies (data download 22 Nov 2021). We highlight in red the positions that are mutated in the 5 current VOCs, as of Dec 2021, indicated in https://cov-lineages.org/index.html: B.1.1.7 Alpha (Panel A), B.1.351 Beta (Panel B), P.1 Gamma (Panel C), B.1.617.2 Delta (Panel D), B.1.1.529 Omicron (Panel E) . We observe a pronounced enrichment for the Omicron variant in the upper right corner, i.e. positions that are likely to mutate (high DCA score), and whose mutations may cause immune escape (high IEDB - RF). Interestingly, the model predicts the positions S371, S373, S375, G496 - mutated in the Omicron variant (Panel E) - to be deleterious, even if they are neutral in the expression experiments (Panel F). As discussed in the main text, this is likely to be due to (a) limited datasets of functional sequences or (b) mutations without effect on expression that may still be deleterious for overall protein fitness. Currently available data are not able to discriminate between these two possibilities.*

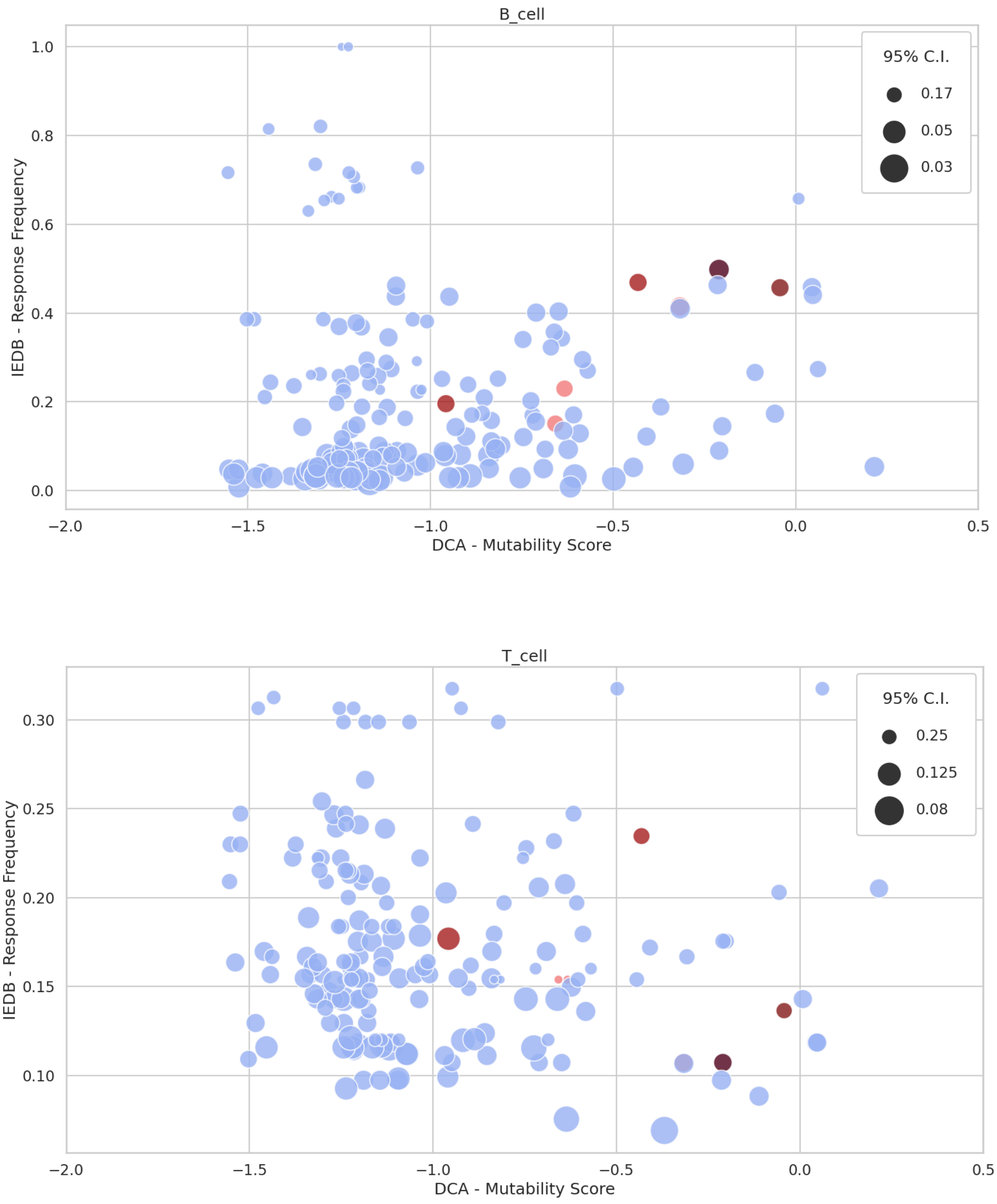


*Fig S8. The IEDB-response frequency considering only B (upper panel) and T (lower panel) cell epitopes, and the DCA mutability score for each position of the RBD domain.*

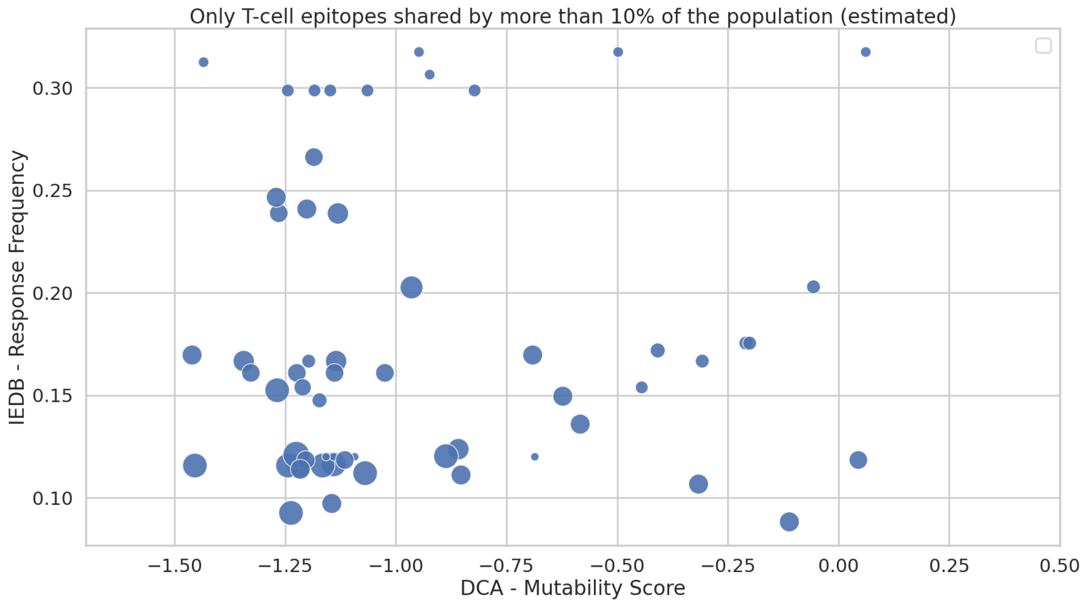


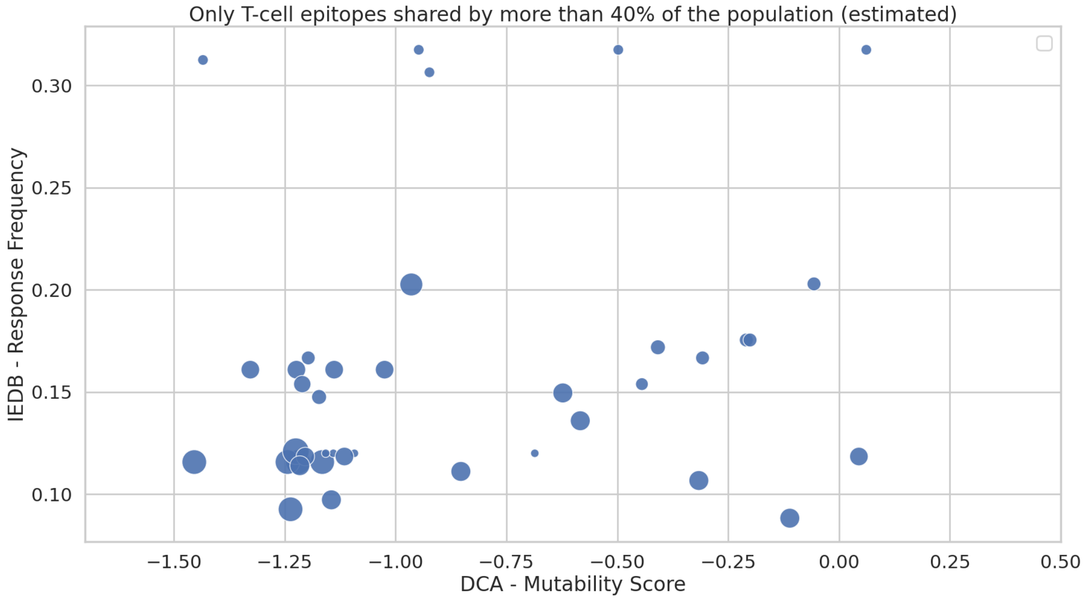


*Fig S9. T cell immunoprevalent epitopes, i.e. predicted to be shared by at least the 10%(upper panel) and 40%(lower panel) of the world population. No clear correlation patterns between DCA and the IEDB emerge with the data available to date. Interestingly, no positions mutated in VOIs/VOCs were identified in T cell immunoprevalent epitopes.*

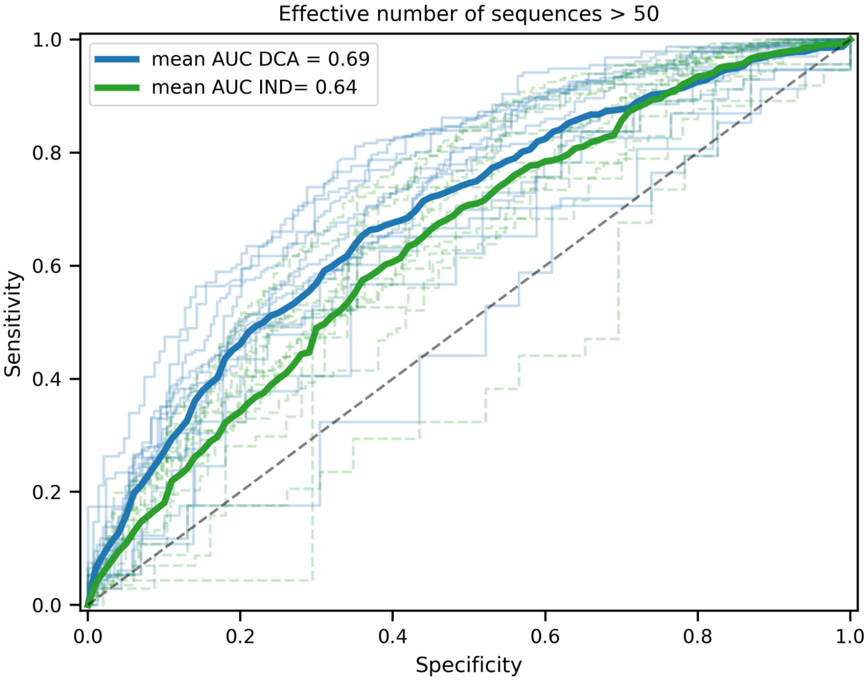


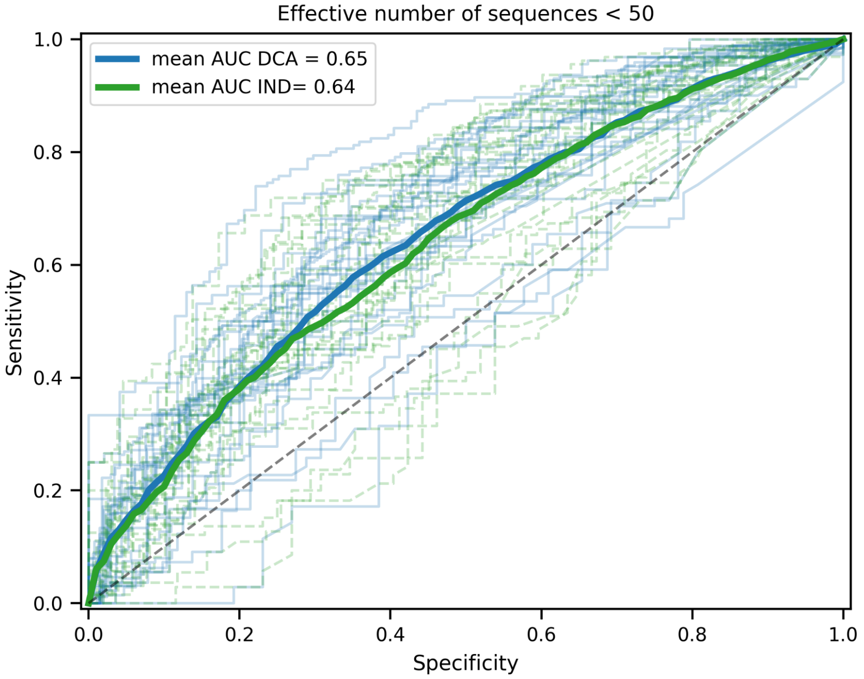


*Fig S10. ROC curve for the DCA (blue) and IND (green) models for all 39 PFAM domains of the SARS-CoV-2 proteome, dividing between (upper panel) domains with more than 50 effective sequences (17 domains, 3491 positions) and (lower panel) less than 50 effective sequences (26 domains, 4546 positions). In bold, the mean ROC curves.*

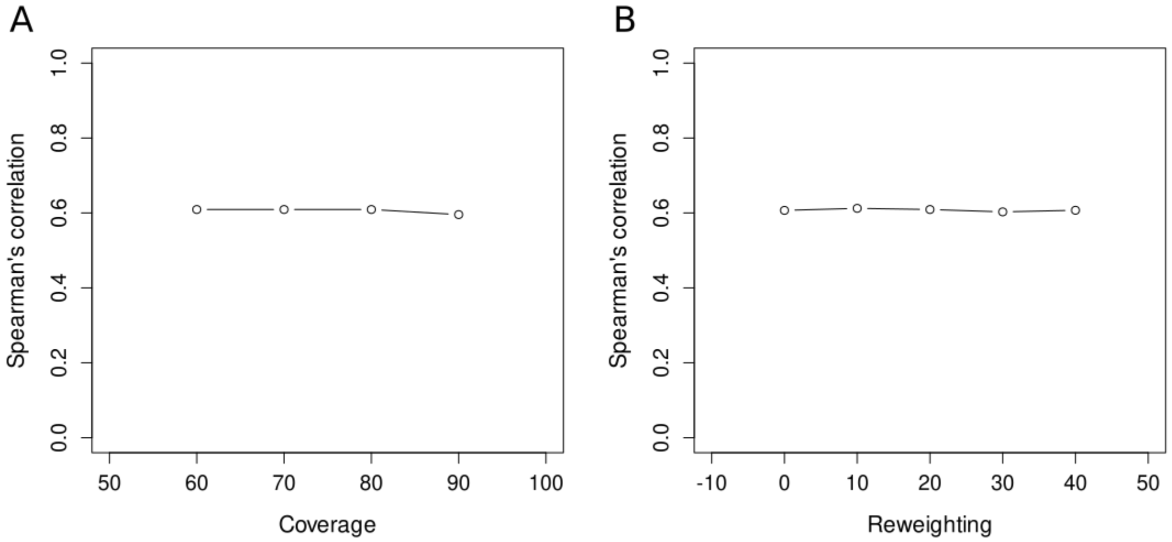


*Fig S11. Spearman's correlation between the DCA model and the observed variability using different thresholds of coverage (filtering out sequences that do not cover that fraction of the reference sequence) (A) or the reweighting parameter (B) for model training.*

*Table S1. List of Pfam protein domains in the SARS-CoV-2 proteome, and the number of effective sequences and positions in the corresponding MSA*

| Protein/ORF | Pfam identifier | Pfam accession | N. eff. seq. | N. positions |
|---|---|---|---|---|
| Envelope | CoV_E | PF02723.15 | 53 | 66 |
| Membrane | CoV_M | PF01635.19 | 40 | 201 |
| Nucleocapsid | CoV_nucleocap | PF00937.19 | 48 | 341 |
| ORF1a | bCoV_NAR | PF16251.6 | 19 | 98 |
| ORF1a | bCoV_NSP1 | PF11501.9 | 12 | 135 |
| ORF1a | bCoV_NSP3_N | PF12379.9 | 9 | 171 |
| ORF1a | bCoV_SUD_C | PF12124.9 | 2 | 64 |
| ORF1a | bCoV_SUD_M | PF11633.9 | 10 | 143 |
| ORF1a | CoV_NSP10 | PF09401.11 | 25 | 123 |
| ORF1a | CoV_NSP2_C | PF19212.1 | 22 | 167 |
| ORF1a | CoV_NSP2_N | PF19211.1 | 25 | 241 |
| ORF1a | CoV_NSP3_C | PF19218.1 | 58 | 488 |
| ORF1a | CoV_NSP4_C | PF16348.6 | 33 | 96 |
| ORF1a | CoV_NSP4_N | PF19217.1 | 58 | 354 |
| ORF1a | CoV_NSP6 | PF19213.1 | 71 | 262 |
| ORF1a | CoV_NSP7 | PF08716.11 | 31 | 83 |
| ORF1a | CoV_NSP8 | PF08717.11 | 30 | 197 |
| ORF1a | CoV_NSP9 | PF08710.11 | 33 | 113 |
| ORF1a | CoV_peptidase | PF08715.11 | 103 | 319 |
| ORF1a | Macro | PF01661.22 | 10075 | 107 |
| ORF1a | Peptidase_C30 | PF05409.14 | 41 | 291 |
| ORF1b | CoV_Methyltr_1 | PF06471.13 | 27 | 522 |
| ORF1b | CoV_Methyltr_2 | PF06460.13 | 31 | 296 |
| ORF1b | CoV_NSP15_C | PF19215.1 | 42 | 153 |
| ORF1b | CoV_NSP15_M | PF19216.1 | 40 | 97 |
| ORF1b | CoV_NSP15_N | PF19219.1 | 38 | 61 |
| ORF1b | CoV_RPol_N | PF06478.14 | 31 | 352 |
| ORF1b | RdRP_1 | PF00680.21 | 61 | 489 |
| ORF1b | Viral_helicase1 | PF01443.19 | 81660 | 225 |
| ORF3a | bCoV_viroporin | PF11289.9 | 3 | 274 |
| ORF6 | bCoV_NS6 | PF12133.9 | 4 | 61 |
| ORF7a | bCoV_NS7A | PF08779.11 | 8 | 106 |
| ORF7b | bCoV_NS7B | PF11395.9 | 3 | 42 |
| ORF8 | bCoV_NS8 | PF12093.9 | 4 | 118 |
| Spike | bCoV_S1_N | PF16451.6 | 103 | 305 |

| | | | | |
|---|---|---|---|---|
| Spike | bCoV_S1_RBD | PF09408.11 | 83 | 178 |
| Spike | CoV_S1_C | PF19209.1 | 6231 | 57 |
| Spike | CoV_S2_C | PF19214.1 | 50 | 40 |
| Spike | CoV_S2 | PF01601.17 | 79 | 522 |

*Table S2. Pfam domains and the number of effective sequences in the MSAs obtained starting with full-length protein sequence or the domain sequence. In bold, the domains where there is a substantial difference.*

| Protein/ORF | Pfam identifier | N. eff. seqs. full-length | N. eff. seqs. domain |
|---|---|---|---|
| Envelope | CoV_E | 53 | 49 |
| Membrane | CoV_M | 40 | 37 |
| Nucleocapsid | CoV_nucleocap | 48 | 47 |
| ORF3a | bCoV_viroporin | 3 | 3 |
| ORF6 | bCoV_NS6 | 4 | 4 |
| ORF7a | bCoV_NS7A | 8 | 8 |
| ORF7b | bCoV_NS7B | 3 | 3 |
| ORF8 | bCoV_NS8 | 4 | 4 |
| **Spike** | **bCoV_S1_N** | **103** | **48** |
| **Spike** | **bCoV_S1_RBD** | **83** | **26** |
| **Spike** | **CoV_S1_C** | **123** | **6231** |
| **Spike** | **CoV_S2_C** | **50** | **7** |
| Spike | CoV_S2 | 78 | 79 |

*Table S3. Strongest inter-domain epistatic couplings for pairs of domains with a maximum (out of all the possible inter-domain epistatic couplings between the pair of domains) coupling higher than 0.5.*

| First protein/ORF | First domain | Second protein/ORF | Second domain | Max. coupling |
|---|---|---|---|---|
| ORF1a | CoV_NSP2_N | ORF3a | bCoV_viroporin | 0.85 |
| ORF3a | bCoV_viroporin | ORF8 | bCoV_NS8 | 0.63 |
| ORF1a | CoV_RPol_N | Spike | bCoV_viroporin | 0.57 |
| ORF1b | CoV_NSP2_N | ORF3a | bCoV_S1_RBD | 0.56 |

### SI text

#### Sequence data

Sequence data in FASTA format were downloaded from the following databases: GISAID (release 16 May 2021), Uniref90 (ref, release December 2020), ViPR (downloaded in September 2020), NCBI viral genomes (downloaded in September 2020) and MERS coronavirus database ( downloaded in September 2020). The amino acid sequence of isolate Wuhan-Hu-1 was used as the reference proteome (genbank identifier: MN908947). Protein domains were detected using the HMMER suite (ref, version 3.1b2) and the HMM profiles from Pfam. After running the command *hmmsearch* from the HMMER suite (ref, version 3.1b2) on the reference proteome using the HMM profiles of SARS-CoV-2 provided by Pfam, the domain amino acid sequence of the full-length protein were trimmed accordingly to the *hmmsearch* output to obtain a reference sequence for each domain. We kept all non-overlapping pfam domains with a domain e-value lower than $10^{-5}$.

A global sequence database including distant species was built by combining Uniref90, ViPR, NCBI viral genomes and MERS coronavirus database. Starting with the domain sequences, we built MSAs by running *jackhmmer* with 5 iterations. For the proteins not belonging to the ORF1ab (which is too long to apply this procedure), we also built MSAs with *jackhmmer* with 5 iterations starting with the full-length reference protein sequences instead of domain sequences. The resulting full-length protein MSAs were decomposed and trimmed to domain alignments by keeping the corresponding columns. When two MSAs from the global database were obtained (one coming from the full-length sequence and another coming from the domain sequence), the one with the highest number of sequences non-redundant at 80% was kept for further analysis for each Pfam domain. Although both strategies usually recover a similar number of sequences, there exists a substantial difference for most domains in the Spike protein (Table S2), allowing us to increase the available sequence data in their MSAs. As quality controls, all sequences including non-standard amino acids were removed as well as repeated sequences or sequences covering less than 80% of the reference domain sequence. To avoid a bias toward the reference sequence, all sequences closer than 90% sequence identity to the Wuhan-Hu-1 reference were filtered out.

For the GISAID database, an MSA for each domain sequence was built using the command *jackhmmer* from the HMMER suite with only 1 iteration as the GISAID sequences are very similar to those in the reference proteome. We filtered sequences including non-standard amino acids, coverage lower than 80%, and those belonging to a non-human host. Only non-identical sequences were considered to avoid the strong sequencing bias due to the highly diverse number of genomes sequenced in different countries. The variability of each position was estimated by counting the number of sequences that have a different amino acid in the corresponding position compared to the reference.

**Co-alignments of domains**

Starting with the 39 raw alignments of domains constructed using both the alignments of distant and close sequences (*Materials and methods*), we build 741 co-alignments (all possible combinations of two domains) by joining the sequences coming from the same genome (thanks to the genome accession number). Co-alignments with fewer than 50 effective sequences were discarded to increase the reliability of the predictions. Note that the number of effective sequences is higher in this analysis compared to the mutability predictions because of the large number of close sequences. From the remaining 601 co-alignments, we computed the models as in case of single domains (see *Material and methods*) and obtained the APC scores from the DCA model between each pair of inter-domains positions. The pairs of domains with at least one APC score higher than 0.3 are linked in Fig. 4D. The 4 predictions with the highest APC scores can be found in Table S3.
